# Neural stimulation hardware for the selective intrafascicular modulation of the vagus nerve

**DOI:** 10.1101/2023.07.14.548991

**Authors:** I. Strauss, F. Agnesi, C. Zinno, A. Giannotti, A. Dushpanova, V. Casieri, D. Terlizzi, F. Bernini, K. Gabisonia, Y. Wu, D. Jiang, V. Paggi, S. Lacour, F. Recchia, A. Demosthenous, V. Lionetti, S. Micera

## Abstract

The neural stimulation of the vagus nerve is able to modulate various functions of the parasympathetic response in different organs. The stimulation of the vagus nerve is a promising approach to treating inflammatory diseases, obesity, diabetes, heart failure, and hypertension. The complexity of the vagus nerve requires highly selective stimulation, allowing the modulation of target-specific organs without side effects. Here, we address this issue by adapting a neural stimulator and developing an intraneural electrode for the particular modulation of the vagus nerve. The neurostimulator parameters such as amplitude, pulse width, and pulse shape were modulated. Single-, and multi-channel stimulation was performed at different amplitudes. For the first time, I polyimide thin-film neural electrode was designed for the specific stimulation of the vagus nerve. In vivo experiments were performed in the adult minipig to validate to elicit electrically evoked action potentials and to modulate physiological functions selectively, validating the selectivity of intraneural stimulation. Electrochemical tests of the electrode and the neurostimulator showed that the stimulation hardware was working correctly. Stimulating the porcine vagus nerve resulted in selective modulation of the vagus nerve. Alpha, beta, and theta waves could be distinguished during single- and multi-channel stimulation. We have shown that the here presented system is able to activate the vagus nerve selectively and can therefore modulate the heart rate, diastolic pressure, and systolic pressure. The here presented system may be used to restore the cardiac loop after denervation by implementing biomimetic stimulation patterns. Presented methods may be used to develop intraneural electrodes adapted for various applications.

## I. INTRODUCTION

The vagus nerve (VN) is the main component of the parasympathetic nervous system and supplies regulatory information to internal organs such as the lungs, heart and most of the digestive system. VN stimulation (VNS) has been exploited extensively to restore the function of the innervated organs. VNS is an FDA-approved treatment for refractory epilepsy and depression that has gathered increased interest in the past decade [1]. Commercial devices are available for VNS, both as implantable (e.g., VNS Therapy®, LivaNova, and Maestro System, EnteroMedics) and transcutaneous (e.g., gammaCoreTM) systems [2]. VNS was proposed as a treatment for various pathologies such as inflammatory diseases (such as psoriatic or rheumatoid arthritis), obesity, diabetes, heart failure, and hypertension [3].There is growing evidence that the VN is involved in the ‘cholinergic anti-inflammatory pathway’, described by the group of Tracey and Pavlov [4]. VNS has been performed with cuff electrodes on 60 patients suffering from chronic heart failure. The therapy showed an improvement in cardiovascular parameters, mainly regarding the left ventricular ejection fraction [5]. However, the mechanism behind this functional recovery is difficult to investigate given the lack of electrodes capable of selectively stimulating cardiac nerve fascicles. Since the VN controls different physiological functions, it is important to be selective when applying neural stimulation. Non-selective neural stimulation may lead to unwanted side effects, especially in the modulation of cardiovascular physiology [6]. Typically, the VN is stimulated using cuff electrodes, which enwrap the epineurium of the nerve, delivering current from its outer layer [1]. Electrical currents applied at the epineurium can lead to non-specific activation of nerve fascicles, leading to adverse side effects [7]. Simultaneous activation of inflammatory, hormonal, or heart-related functions may result from non-selective stimulation. Alternative solutions to commercial cuff electrodes have been proposed to increase the selectivity of the stimulation. For example, cuff electrodes with multiple contacts distributed radially on the nerve, microwires penetrating the nerve, or intraneural electrodes [8]. VNS allowed to modulate the heart rate (HR) and the blood pressure (BP) and treat hypertension in rats by stimulating the left vagus nerve (LVN) [9]. Microwires and intraneural electrodes are inserted into the nerve crossing the epineurium. Microwires and intraneural electrodes are inserted into the nerve crossing the epineurium which acts as an electrical insulator. Electrodes inserted into the nerve overcome this barrier. Therefore, smaller currents are sufficient to elicit axons. Furthermore, neural selectivity is increased due to smaller electrical fields created during neurostimulation. There are different types of intraneural electrodes, that can be inserted longitudinally (longitudinal intrafascicular electrode, short LIFE) or transversally (transversal intraneural multi-channel electrode, short TIME) into the target nerve. Both LIFE and TIME have been used to restore sensory feedback in amputees with promising results, demonstrating stability of intraneural stimulation up to 6 months [10]. In 2022, Jayaprakash et al. have shown in the domestic porcine model that fibers cluster across the length of the vagus nerve according to their function [2]. This type of fascicular organization suggests that a transversal arrangement of small, intraneural stimulation sites could be ideal for VNS applications. However, experimental evidences assessed in clinically relevant models to support this suggestive hypothesis are ongoing. The goal of our study was to develop neurostimulation hardware specifically tailored for the selective stimulation of the cardiac fascicles of the VN. The function of the neurostimulation hardware should be validated by the reproduction of previously shown modulation capacity of the VN [9]. For this purpose, we developed an intraneural thin-film electrode (IE) based on the TIME concept, adapted an existing handheld neurostimulator for use with the developed electrode and developed a graphical user interface (GUI) to control the neurostimulator. It is the first time that previously successful polyimide thin-film electrodes are used to stimulate the VN. The IE was designed based on the dimensions of histological sections of porcine VNs to best target the distribution of active sites (AS) and match its specific morphology. In fact, the human and porcine VNs have comparable size, similar amounts of fibrous tissue and contain multiple fascicles. The IE was characterized electrochemically *in vitro* following established procedures. *In vivo* proof of concept experiments were performed recording electrically evoked action potentials (eCAPs), BP and HR during intraneural stimulation.

## II. METHODS

### A. Electrode design specifications

The development of the IE followed previously suggested guidelines [11]. The following approach was used: 1) Obtaining histological transections of the porcine VN. 2) Development of an IE, based on histological samples, to guarantee optimal AS distribution and therefore neural selectivity. 3) Electrochemical, and structural characterization of the electrode to verify its functionality and limitations in terms of maximum injectable charge (MQ_inj_). 4) Adaptation of an existing neurostimulator to the IE performance. Development of a GUI to control intraneural stimulation. 5) In vivo testing to validate the neural selectivity of the developed stimulation hardware. The workflow is highlighted in **Figure 1, A to F**. The amplitude modulation of eCAPs and the changes observed in the physiological parameters following neural stimulation should confirm that the here developed neural stimulation hardware allows for the selective stimulation of the VN.

**Figure 1.**
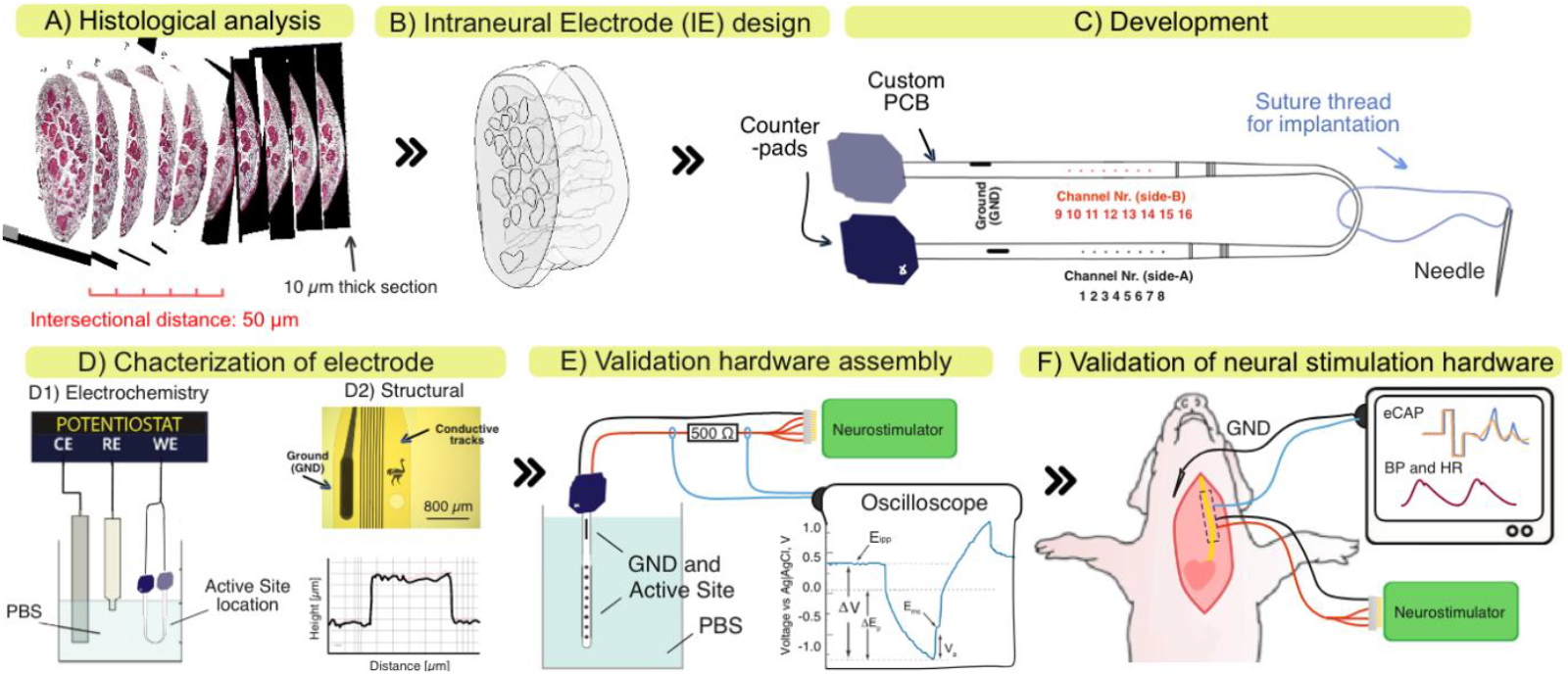
Workflow for the development and validation of neural stimulation hardware to selectively modulate the vagus nerve. A) Histological transections of the vagus nerve (VN). B) 3D nerve model of explanted VN. C) Features of developed of intraneural electrode (IE). Assembly with custom-made PCB used to connect the intraneural electrode to the neurostimulator. D1) Electrochemical and D2) structural characterization of the IE. Electroscopic Impedance Spectroscopy (EIS) and Cyclic Voltammetry (CV) measurements were performed. E) Validation of neural stimulation hardware assembly to evaluate maximum injectable charge (MQinj). F) Stimulation of VN and recording of electrically evoked potentials (eCAP), heart rate (HR), diastolic pressure (DP), and systolic pressure (SP) to validate selectivity of the intraneural electrode (IE). Figure adapted from [8].

#### 1) Histological analysis of VN

Two male, healthy, sexually mature Göttingen minipig (Ellegaard Göttingen Minipigs A/S, Dalmose, Denmark; avg body weight 35 kg) were enrolled in this first feasibility study. One animal was used for histological analysis. The other was used for the stimulation experiments. The Animal Care Committee of the Italian Ministry of Health approved this study, which was conducted in accordance with Legislative Decree No. 26/2014 (2010/63/EU). Histological transections of one LVN and one RVN of the healthy animal were obtained. The explantation location was at the mid-level of the cricoid. The nerves were cut through at the distal level using a scalpel and fixated onto a styrofoam support in an elongated state to prevent retraction of the tissue. The nerve samples, including the styrofoam support, were immersed in 15 ml falcons filled with formalin 4 % and stored in a fridge at 4-6 C° for 20 hours. Next, the nerves were washed three times in room temperature standard 1% phosphate buffer saline solution (PBS) for 15 min each and stored in 70 % ethanol at 4-6 C° until further processing. The proximal end was cut obliquely, while the distal end was cut transversally to identify the nerve direction after the next steps. The nerves were placed in histological sample cassettes and immersed in an ethanol solution of 80%, 90%, and 100% concentration for one hour each for dehydration. Ethanol was substituted with Xylol by immersing the samples twice in 100% Xylol for 60 minutes each. The nerves were then immersed and maintained in Paraffin for 6–12 hours. The nerves were then embedded completely with Paraffin wax (HistoCore Arcadia H and C, Laica, US), cooled down, and transected using a microtome (Leica SM2010R, USA). Every 50 μm, 10 μm thick transversal sections were cut, and mounted onto microscope glass slides. The transections were stained with Hematosin and Eosin by immersion. Images were taken (Leica DMi8, GER) of each transection to create a 3D model in the computer aided design (CAD) software Solidworks (Dassault, FR). The histological transections and the 3D model of both VNs can be seen in **Figure 1, A and B** respectively. Materials described in this section were acquired from Sigma-Aldrich (US) if not stated otherwise. From the 3D model, the maximum and minimum nerve diameter, the number of fascicles, the nerve section area, and the fascicle area were measured for each section. The average (avg) and standard deviation (std) were calculated for each parameter and used to design the IE to maximize neural selectivity.

#### 2) Electrode design and development

The electrode design consisted of a foldable polyimide structure which was inserted into the nerve through a suture thread attached to the tip of the IE (**Figure 1, C)**. We adapted the AS number, and location based on the morphology of the porcine 3D models to place as many AS as possible within the nerve itself. Eight AS (ø = 80 μm) were placed on each side of the electrode for a total of 16 ASs. To maximise the number of AS the inter-AS distance was reduced from 750 μm (TIME) to 400 μm. AS should be equally distributed on the inside of the nerve after implantation. The electrode consisted of a 10 μm thick polyimide sandwich structure developed with photolithographic methods. Here, we shortly describe the photolithographic development process. A sacrificial layer of Ti/Al (20/100 nm) was deposited on a 4-inch silicon wafer by e-beam evaporation. Next, a 5 μm-thick layer of polyimide (PI2611, HD Microsystems GmbH, GER) was spin-coated on the wafer and cured for 2 hours at 300 °C in a N_2_ oven for hard bake. The first photolithographic step was performed by spin-coating, baking, exposing, and developing a 4 μm-thick photoresist (ECI 3027, MicroChemicals, GER). As conductive film, Ti/Pt/IrOx (25/300/400 nm) was sputtered (AC450, Alliance Concept) and patterned through lift-off according to previous literature [8]. The AS material in contact with the nervous tissue was coated with IrOx, therefore sputtered IrOx was applied (SIROF). Next, a second 5 μm-thick layer of polyimide was spin-coated and cured to encapsulate the tracks. A second photolithography was performed using a 12 μm photoresist (AZ10XT, MicroChemicals, GER), followed by O_2_-based reactive ion etching to shape the IE and expose the electrodes, AS, and contact pads. Finally, the individual devices were released through the anodic dissolution of aluminum in a 1.5 V bias in saturated NaCl solution. To connect the IE to the neurostimulator a custom-made printed circuit board (PCB) was developed (**Figure 1, C)**. The exposed pads of the basic electrode structure were soldered to the PCB using the silver conductive glue Ablestick 84-1LMI (Henkel, GER). Roughly 0.1 g of glue was deposited onto the exposed pads using a pressure dispensing system (EFD, US). The polyimide structure was aligned with the PCB, and connected through the application of light pressure onto the polyimide structures. Next, the assembled structure (IE including PCB) was baked at 120 C° for 120 minutes until the silver glue was cured. 100 μm thick copper wires (RS components, ITA) were soldered to each counter pad of the PCB (**Figure 1, C)**. Counter pads were connected to a male connector A79022001 (Omnetics, US) by manual soldering. The mapping procedure of each AS channel can be seen in **Figure 1, B**. The AS furthest closest to the PCB was considered to be AS1. Since each side had eight contacts, side A was mapped to be AS1-8, and side B was mapped to be AS9-16. Finally, the PCB was insulated to prevent current leakage. We applied the UV-curable, biocompatible glue 1401-M-UR (Dymax, GER) to cover the IE and PCB. The glue was cured, by applying 60 seconds of UV light. Finally, a suture thread with a loop on one side and a needle on the other (EH7900G Prolene, Ethicon, US) was secured to the electrode as shown in **Figure 1, C** to allow insertion of the electrode in the nerve.

#### 3) Electrode characterization

An electrochemical and structural analysis was performed to validate the photholitographic process (**Figure 1, D1 and D2 respectively)**. A visual inspection was performed using a microscope (Hirox, JAP) to check for interruptions in the conducting tracks of the electrodes. Electrodes with interruptions were discarded. The same method was used to measure the real AS diameter, necessary to calculate the real cathodic charge storage capacity (cCSC) [8]. The real AS diameter was compared with the expected value of 80 μm. To characterize the precision of the photolithographic process, the AS metallization height (ASH), the track metallization height (TH), and the height of the final polyimide structure (PH) were measured using a mechanical profilometer (Dektak XT, Bruker, US). ASH and TH were measured after the patterning of the conductive Ti/Pt/IrOx film. The PH was measured after RIE and before lifting off the electrodes from the wafer. The ASH, TH, and PH were obtained from the profilometry data using a Matlab (Math Works, US) based software. Averages, and standard deviations were calculated. Results were compared with the design values. A comparison between IE from the most lateral and the central location of the wafer substrate was performed. Structural results were used to validate the repeatability and precision of the lithographic development process. To validate the electrical functionality, electrochemical impedance spectroscopy (EIS), and cyclic voltammetry measurements (CV) were performed on eight ASs using a potentiostat (Gamry Instrumments, US). Both measurements were performed in a standard 1% phosphate buffer solution (PBS). Shortly, a large Pt mesh counter electrode, an Ag/AgCl reference electrode, and the IE were immersed into the PBS solution (**Figure 1, D1**). During EIS, 10 Hz to 100 kHz RMS, 10V p.p. AC waves were delivered through each AS to measure the impedance. The avg impedance and std were calculated for all active sites at a physiological reasonable frequency of 1 kHz. The resulting phase was analyzed for resistive or capacitive behavior. We compared impedance to previously reported values for similar electrodes. CV should give indications on the reversible of reactions such as oxidation of the electrode surface during stimulation. Therefore, in the same setup, 20 CV cycles were performed for each AS to measure the cathodic charge storage capacity (cCSC) in vitro. A water window for IrOx of -0.6 V to +0.8 V was defined according to Cogan et al. 2009 [12]. The cCSC gave information about the maximum charge which could be injected through each AS without causing damage through oxidation or reduction processes. The time integral of the cathodic current was calculated to evaluate the cCSC for the electrodes AS. Cycling was performed with a scan rate of 50 mV/s, between -0.6 and 0.8V as potential limits in PBS [8].

### B. Neurostimulator

#### 1) Hardware and stimulation performance

To drive the electrodes, a handheld 16 channels, neurostimulator was designed and tested in combination with the IE (**Figure 1, E)** before performing the *in vivo* experiments (**Figure 1, F**). The Neurostimulator consisted of an integrated circuit stimulator (0.18 μm CMOS high-voltage technology) and a peripheral block, controlling the integrated circuit stimulator. The peripheral block provided power and digital commands sent by the user through a GUI (**Figure 2, A and B)**. The Neurostimulator was powered through a micro USB cable attached to a PC running on battery. The stimulator chip was mounted into a 48-pin quad flat no-lead package. The die area was 1960 μm x 6100 μm. The package was assembled with a PCB containing the peripheral block components. To connect with the 16 ASs of the IE contacts, a 32 channel A79025 connector (Omnetics, US) was mounted to the front of the Neurostimulator (**Figure 2, C)**. Biphasic stimulation current was implemented to maintain electrochemical balance at the electrode tissue interface. The injection of bi-phasic, square waves should reduce oxidation or reduction reactions, reducing the electrodes current injection capacity [8]. Each channel of the Neurostimulator could independently provide up to 500 μA of rectangle, charge-balanced, biphasic, cathodic-first stimulation current. The voltage compliance was 20 V requiring 23.3 μW when in standby mode. When stimulating, the compliance is dynamically based on the imposed stimulation pattern. With passive discharge, at a 100 Hz asymmetrical pulse rate, the measured residual DC current was 3.9 nA which was well below the 100 nA recommended safety limit [13]. During the characterization the stimulator was connected to an RC electrode model (Rx=100k, Cx=100nF parallel, then in serial with Ri=2k) for the test. The biphasic current pulse parameters were set as follows: cathodic current 200 μA, pulse width 100 μs, interphase delay 25 μs, anodic current 50 μA, pulse width 400 μs. The pulse frequency was set to 100 Hz, which is a more strict condition for charge balancing. A passive discharge period of 450 μs is inserted after each completed biphasic cycle. Residual current was measured on another serial resistor of 100R, by measuring the voltage and dividing R. The Neurostimulator was previously developed and adapted for the current study. Current waveforms for each channel during multichannel stimulation should were recorded and verified. The goal was to provide current patterns which have previously been reported with similar intraneural electrodes used in human peripheral nerves [10]. Possible stimulation parameters are described in **Table 1**. Anodic current and pulse width were automatically calculated by setting the anodic ratio according to the stimulation charge net-zero principle. The maximum anodic width allowed was 2000 μs. The neurostimulators code was designed to check the values prior to stimulation. An FPGA Cmod-S-7 module inside the Neurostimulator provided the digital control and communicated signals via a universal asynchronous receiver-transmitter (UART) interface between the PC and the stimulator chip. Two instrumentation amplifiers (IA) were incorporated to monitor the simulation voltage pattern (Ve) on the selected electrode and the output biphasic current pulses by measuring the voltage (Vi) across a 500 Ω sensing resistor shown in **Figure 2, D**.

**Table 1.** Stimulation range and step size for parameters which can be imposed for the Neurostimulator.

|  | Cathodic current | Cathodic pulse-width | Anodic ratio | Pulse interphase | Pulse intervals |
| --- | --- | --- | --- | --- | --- |
| Range | 0-500 μA | 20 – 2000 μs | 1 – 10 | 100 - 100 μs | 10 – 1300 ms |
| Step-size | 1 μA | 10 μs | 1 | 10 μs | 1 ms |

**Figure 2.**
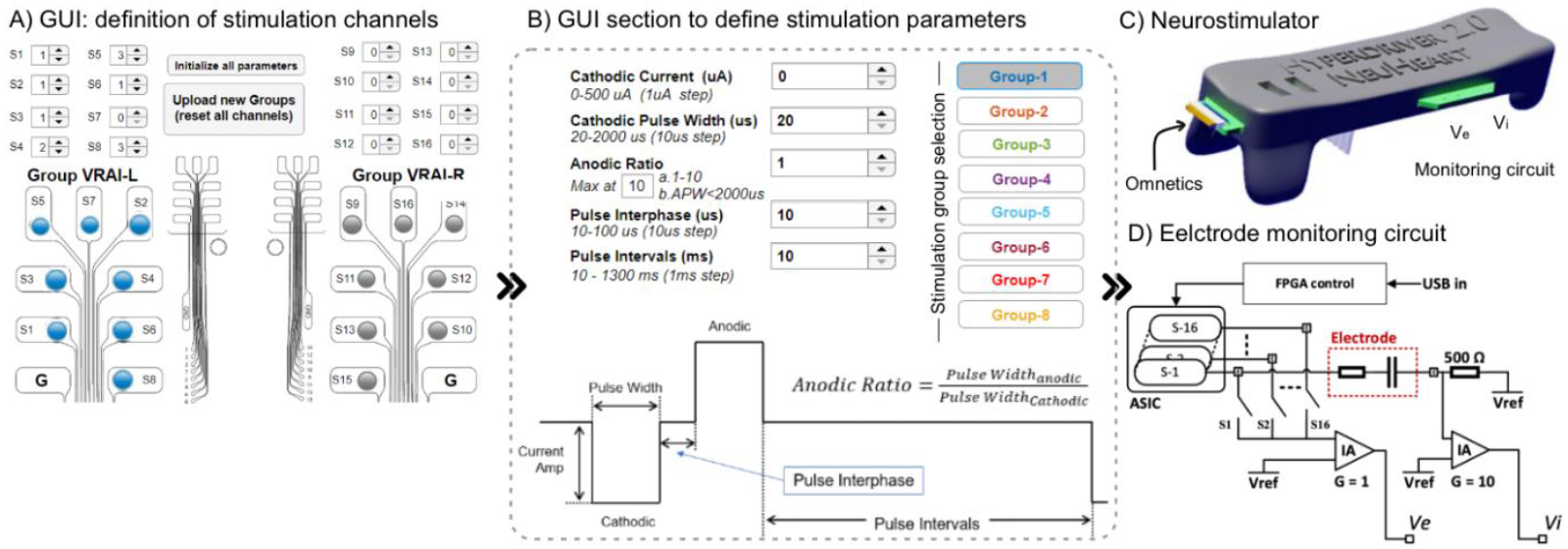
A) The graphical user interface (GUI). Single-channel and multi-channel stimulation could be defined for up to 16 Channels. Each stimulation channel operated individually. Gray channels were not active. B) Definition of the stimulation parameters. Current amplitude, pulse width, anodic ratio, pulse interphase, and pulse interval could be defined. C) Image of the neurostimulator and the monitoring circuit. D) The electrode monitoring circuit. Ve gave information about the voltage on the selected active contact. The 500 Ω resistor in the return path was used to measure the current going through the electrodes.

#### 2) Neurostimulator control platform

In previous publications, the use of a GUI to control the neurostimulation platform has shown to be very useful during *in vivo* experiments. Therefore, we developed a GUI (**Figure 2, A & B)**, implemented in Matlab, which allowed to perform single-channel or multi-channel stimulation of specific channels. During single channel stimulation, one of 16 AS could be stimulated with the parameters imposed through the GUI. AS were defined as S1-S16. During multi-channel stimulation, the stimulation parameters were sent through two or more channels at the same time. AS were defined as groups (**Figure 2, A & B)**. During the present study one group of 8 AS was defined, as can be seen in **Figure 2, A**. The electrode mapping was shown in the middle of the GUI. The stimulation parameters were selected through the tab “Channel Parameters” shown in **Figure 2, B**.

#### 3) Testing the neurostimulator and VNS

The validation of the hardware equipment prior to animal experiments was of great importance to assess functionality. Failures in experimental setups during animal experiments may lead to the abandonment of the experiment, and the unnecessary loss of the animal. To increase the chances of success, and reduce animal use in experiments, we tested our experimental setup for *in vivo* stimulation in the laboratory. To validate the electrical functionality of the neurostimulator together with the electrode, voltage transient measurements (VTm) were performed for each channel. The electrode was immersed in PBS and connected to the neurostimulator (**Figure 1, D1)**. Each channel of the neurostimulator was connected to one channel of the electrode. The neurostimulator was connected to a battery-powered PC running the GUI. The return electrode (GND) of the neurostimulator was connected to the GND of the electrode. An oscilloscope (DS1102E, Rigol, CN) was attached to the monitoring circuit pins of the neurostimulator (**Figure 1, E** and **Figure 2, C)**. We measured the voltage drop over the 500 Ω resistor. Rectangle, charge balanced, bi-phasic, cathodic-first currents were injected through the electrode’s AS. The injected currents ranged from 50 μA to 500 μA in steps of 50 μA and reflected the currents which should be injected during *in vivo* experiments. The pulse-width was fixed at 200 μs. The frequency was set to 5 Hz. The resulting VTm were acquired using the oscilloscope. The water window was defined to be in a range of -0.6 V to +0.8V. Using a Matlab function, the maximum cathodically driven electrochemical potential excursion (E_mc_) was calculated for each AS and current amplitude. E_mc_ was defined according to Cisnal 2019 et al. [14]. Shortly, E_mc_ was defined as the sum of the interpulse potential E_ipp_ and the polarization across the electrode-electrolyte interface E_p_. The polarization was further defined as the difference of the overall voltage drop ΔV and the access voltage V_a_. Graphical information on how the mentioned parameters were defined can be found in **Figure 1, E**.

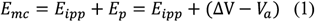

*V*_*a*_ was evaluated, calculating the deviation of the VTm. *V*_*a*_ was the voltage value of the VTm measured at the end of the first cathodic pulse injected during VTm. To evaluate the maximum injectable charge, the injected current at the highest E_mc,_ which was not bigger than 0.6 V, was considered. The resulting E_mc_, maximum injectable current (IC_max_), and maximum injectable charges were averaged over eight AS. MQ_inj_ were calculated by multiplying the injected current amplitude, and the pulse-width.

### C. Experimental setup

#### 1) Animal preparation

The second minipig was used for the stimulation experiments. The animal was premedicated with Zoletl® (10 mg/kg), anesthetized with Propofol (2 mg/kg intravenously) and maintained under 1-2 % sevoflurane in air enriched by 50 % oxygen. The animal received an infusion of 500 mL NaCl (0.9 %) solution to prevent dehydration. The animal was placed and kept in a dorsal position [15]. The experimental setup is shown in **Figure 1, F**. The legs were fixed with velcro fasteners to the surgery table to assure stability. Oxygen saturation, arterial pressure, and heart rate were continuously monitored. To expose the thoracic part of both VNs a sternotomy was performed. Blunt dissection was used to expose the RVN both at the cervical and thoracic level. The VN were surgically exposed by removing surrounding connective tissue. The VN was lifted distally and proximally using two surgical latex rubber bands, held in position using forceps locking scissors. The nerve was held in a slightly tense position, which facilitated the insertion of the electrode. One IE was implanted to perform intraneural stimulation. To fixate the electrode, KWIK-CAST (World Precision Instruments, US) was applied to the insertion location. A 6 channel cuff electrode (Microprobes, US) was positioned 1 cm rostrally with respect to the IE to record eCAPs. After implantation, the latex rubber bands were released and the nerve turned to a relaxed position. The implanted IE and cuff can be seen in **Figure 3, A**. The surgical opening was filled with sterile physiological solution to guarantee that the GND of the IE was in contact with the animal and not floating in the air. The IE and the cuff were connected to the TDT (Tucker Davis Technologies, US). Recording, stimulation, and impedance measurement were performed to confirm the correct experimental setup. Subsequently, the electrode was connected to the neurostimulator. The cuff electrode was connected to a PZ5 neuro digitalizer and a RZ5 BioAmp processor to record eCAPs.

**Figure 3.**
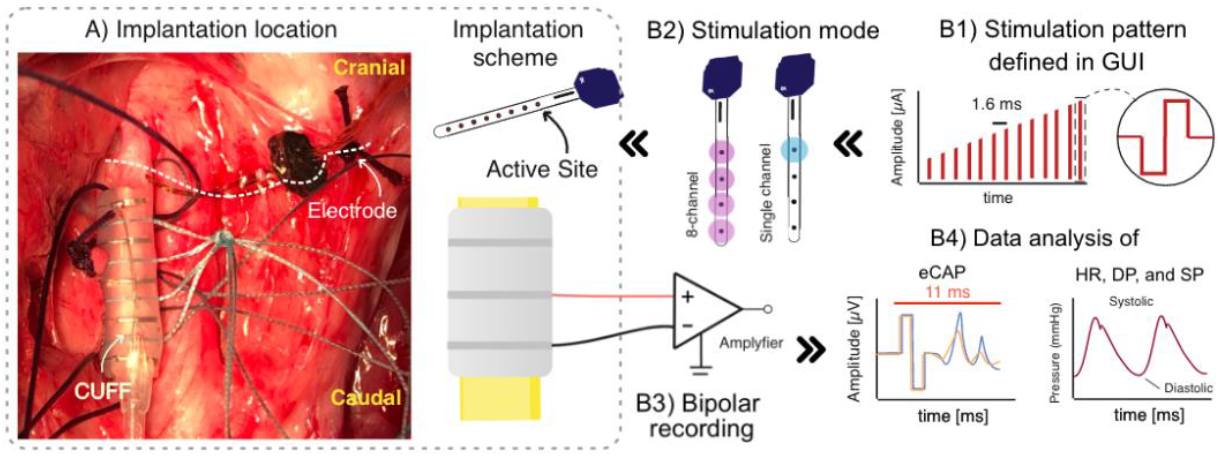
Overview of the experimental setup to validate the selectivity of neural stimulation hardware. A) Implantation location of intraneural electrode and CUFF electrode. Of five CUFF contacts, only three were used. Not seen on the image: the implantation location was filled with saline solution to guarantee that the intraneural electrode GND is in contact with the animal. B1) Stimulation patterns were defined in the GUI used to control the neurostimulator. The pulse width was fixed at 250 μs, the frequency was fixed at 100 Hz. The current amplitude raged between 100-500 μA. B2) Intraneural contacts were stimulated singularly or in multiple configurations (one or eight channels respectively). B3) The eCAPs were recorded using a bipolar configuration. B4) The data were analyzed offline, averaging the resulting eCAPs and physiological data. The analysis window for the eCAPs was 11 ms and started 1 ms before the stimulation trigger onset.

#### 2) Validation of selective stimulation

To evoke vagal eCAPs in the RVN rectangle, bi-phasic, charge balanced, cathodic-first pulses were injected. The current amplitude ranged from 100 to 500 μA, the pulse-width was fixed at 200 μs, with a frequency of 30 Hz and an interphase interval of 10 μs (**Figure 3, B1)**. Two types of stimulations were performed. Either one (single-channel) or eight AS (multi-channel) were stimulated (**Figure 3, B2)**. As return electrode, the GND of the electrode was used. The PZ5 was set to bipolar recording mode. As GND electrode for the eCAP recording, a 1.0*15 mm tungsten needle was inserted subcutaneously, in the most superficial muscular tissue of the surgical opening (**Figure 1, F)**. Neural recordings were sampled at 25 KHz and saved as raw data by the RZ5 BioAmp software Synapse. The recording was started 30 seconds prior to stimulation to obtain baseline reference. During stimulation experiments, the heart rate (HR), diastolic pressure (DP), and systolic pressure (SP) were recorded to evaluate if intraneural stimulation had effects on the modulation of physiological parameters (**Figure 3, B4)**.

### D. Data analysis

Data processing and analysis was performed using Matlab. *Neural recordings:* The electroneurograpy data were filtered using a 4th order butterworth 50 Hz highpass filter. The timing of each stimulation pulse was identified through the respective stimulation artifact. Stimulation triggered averaging was used to evaluate the presence of eCAPs. The analysis window was 11 ms large, and started 1 ms before the stimulation trigger was recorded (**Figure 3, B4)**. Systolic and diastolic arterial blood pressure (SBP and DBP, respectively) were calculated for each heart beat and filtered to remove the variation induced by the breathing cycle. *Physiological data analysis:* The heart rate (HR) was derived from the arterial BP signal, inverting the time interval between each subsequent SBP peak. DBP and SBP were calculated and plotted against time.

## III. RESULTS

### A. Electrode design based on VN histology

Transections of one LVN and one RVN were obtained, performing histological analysis. Ten and six, 10 μm thick sections were obtained from the LVN and RVN respectively. The distance between each section was 50 μm. Therefore, a total length of 0.5 mm (LVN) and 0.3 mm (RVN) was analyzed. All results regarding the histological analysis are shown in **Table 2**. The average maximum diameter of the LVN sections was 2.6 mm (std +/- 0.1) corresponding to a variation of 3.9 % between sections. The average maximum diameter of the six RVN sections was 2.8 mm (std +/- 0.03) corresponding to a variation of 1.1 % between sections. The average minimum diameter for the LVN was 2 mm (std +/- 0.08) corresponding to a variation of 4.1 % between sections. For the RVN the average minimum diameter was 1.9 mm (std +/- 0.04) corresponding to a variation of 2.1 %. We counted the number of fascicles for each transection. Within the LVN and RVN, 26.9 (std +/- 1.79) and 37.2 (std +/- 1.94) fascicles were counted respectively. The average area of the transections was 4 *mm*^2^ (std +/- 0.09) and 4.3 *mm*^2^ (std +/- 0.13) for the LVN and RVN respectively. Hence, the RVN showed a higher surface area. The fascicle area was measured to be 1.2 *mm*^2^ (std +/- 0.36) and 1.6 *mm*^2^ (std +/- 0.22) for the LVN and the RVN respectively. The fascicle area of the LVN was therefore larger compared to the RVN.

**Table 2.**
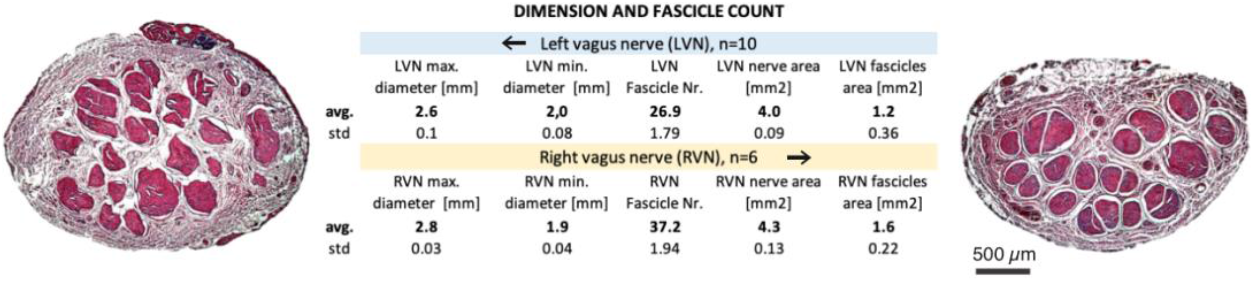
Results of histological analysis of one left and one right VN sample. The total length of the analyzed sample was 300 μm and 500 μm for the LVN and RVN respectively. Ten and six slices were analyzed for the LVN and RVN respectively. The left image shows section six of the LVN. The right image shows section six of the RVN. The scale bar is valid for both images.

**Table 3.**
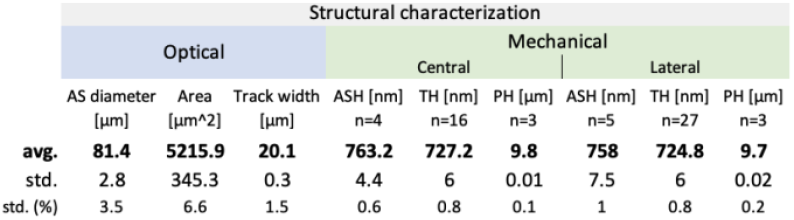
Results of structural characterization of five central, and six lateral AS. The active site height (ASH), Track height (TH), and height of the polyimide structure (PH) were validated. The PH corresponds to the intraneural electrode (IE) device height.

### B. Electrode characterization

#### 1) Photolithographic development process

The average real AS diameter (**Figure 4, A**) was measured for 16 ASs to be 81.4 μm (std +/- 2.8) compared to 80 μm expected diameter. The average surface of the real AS was 5215.9 μ*m*^2^ (std +/- 345.3) compared to the expected 5026 μ*m*^2^. The track width was measured in 11 locations and resulted to be 20.1 μm (std +/- 0.3) compared to 20 μm expected diameter. All results of the optical characterization measurements can be seen in **Table 2**. The ASH, TH, and PH were measured to characterize the precision of the photolithographic development process. The ASH was measured for four center AS and five lateral AS and resulted to be 763.2 nm (std +/- 4.4) and 758 nm (std +/- 7.5) respectively for an average difference of 5.2 nm between the center and lateral electrodes. An average ASH height of 760 nm was measured. The expected height was 725 nm (Ti/Pt/IrOx). Therefore, a difference of 35 nm was measured. The TH was measured for 16 center electrodes, and 27 lateral electrodes and resulted to be 727.2 nm (std +/- 6) and 724.8 nm (std +/- 6) respectively. A difference of 2.4 nm between center and lateral IE was the result. The average of the TH was 725.7 nm. The PH was measured for three lateral and three center IE. The PH resulted to be 9.8 μm (std +/- 0.01) and 9,7 μm (std +/- 0.02) for the center and lateral IE respectively. A difference of 0.1 μm was measured between the center and lateral electrodes. Compared to the expected height of 10 μm, we measured a difference of 0.25 μm. All results of the characterization measurements can be seen in **Table 2**.

**Figure 4.**
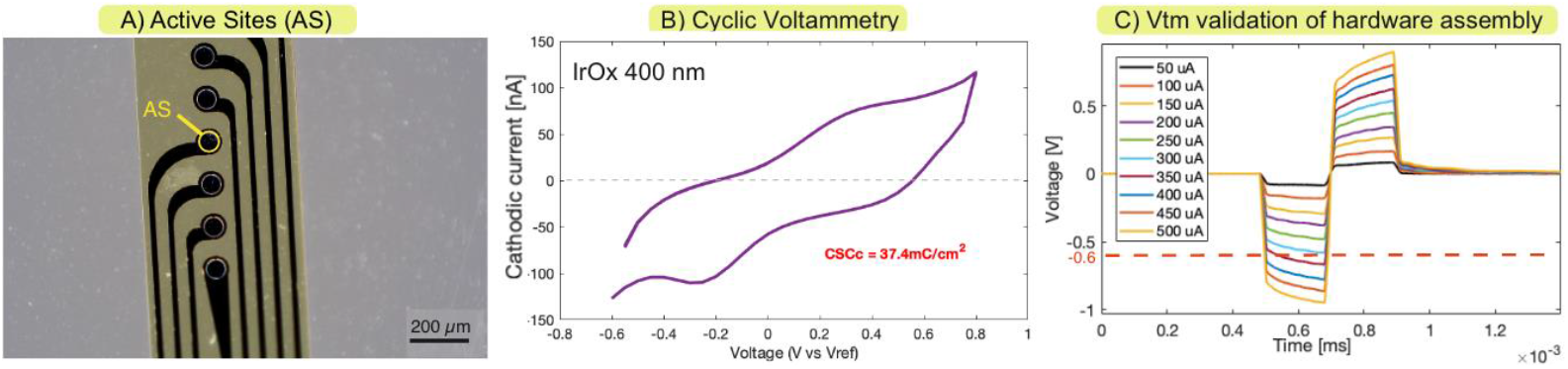
A) Optical picture of one side of the intraneural. The electrodes were checked for interruptions in the conductive tracks and measuring the AS surface. The AS surface was further used to calculate the cCSC. B) One of twenty Cyclic Voltammetry (CV) of IrOx coated AS. C) Voltage Transient measurement example to validate if the maximum current of the neurostimulator exceeds the water window of the intraneural electrode. Emc is indicated for AS two.

#### 2) Electrochemical studies

EIS, CV, and VTm measurements were performed on 8 ASs using the standard three-electrode setup. EIS measurements of IrOx coated AS performed at 1 kHz showed an average impedance of 12.4 kΩ for 16 ASs (std +/- 5). As reported previously, EIS starts to stabilize between 100 Hz and 1 kHz in terms of pitch. The phase behavior was confirmed to be resistive, as has been previously reported for IrOx [12]. The standard deviation was quite high at 40.3 %. The impedance compared between IE sides (AS 1-8 and 9-16) were 8.5 kΩ (std +/- 1.8) and 15.8 kΩ (std +/- 4.5) respectively. Results are shown in **Table 4**. CV was performed to calculate the cCSC. Calculations were based on the measured average AS surface of 5215.9 μ*m*^2^ and resulted to be 37.4 *mC*/*cm*^2^ (std +/- 7.1). An example of a typical CV curve is shown in **Figure 4, B**. Finally, a counterclockwise inclination can be observed in the CV plot. Results are shown in **Table 4**. VTm measurements were performed in PBS using the neurostimulator and oscilloscope. The water window we considered was - 0.6 V for the cathodical pulse, is indicated by a red line. E_mc_ within the water windows were measured to be 0.52 mV (std +/- 0.7). IC_max_ resulted to be an average of 492.9 μA (std +/- 18.9). VTm resulted in an average MQ_inj_ of 98.6 *nC* (std +/- 3.8) per AS. For all AS it was possible to stimulate with 500 μA without reaching the water window. Only AS one reached the water window at a IC_max_ of 450 μA. The MQ_inj_ for AS two was 90 nC. For all other AS a MQinj of 100 nC was measured. All AS were quite close to the water window, with an average E_mc_ of 0.52. An example of the VTm for AS 2 can be seen in **Figure 4, C**. E_mc_ at 0.54 V is indicated at a current intensity of 500 μA. The results of the electrochemical analysis highlighting the functionality of the electrode and the neurostimulator are shown in **Table 4**.

**Table 4.**
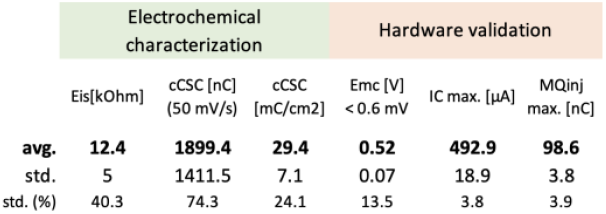
Data obtained performing electrochemical impedance spectroscopy (EIS), cyclic voltammetry (CV) of eight intraneural electrode AS. The calculations of cCSC were made based respecting the dimension of each AS.

### C. eCAP recording from the right vagus nerve

A comparison between single-channel and multi-channel AS stimulation and resulting eCAPs was performed (**Figure 5, A**). The stimulation delivered activating all eight AS produced a significantly larger evoked response. The eCAPs recorded during stimulation delivered using eight AS were characterized by a first positive peak (P1) at 1.04 ms with an amplitude of 59.4 μV, a first negative peak (N1) at 1.16 ms with an amplitude of -12.5 μV, a second positive peak (P2) at 1.38 ms with an amplitude of 23.6 μV, a second negative peak (N2) at 1.79 ms and an amplitude of -20.5 μV, and a third positive deflection at 3.5 ms and an amplitude of 5.8 μV. In contrast, the eCAPs recorded during stimulation delivered using a single AS did not have a clearly visible P1, N1, and P3, while P2 and N2 had amplitudes of 8.8 μV and -7.3 μV. eCAPs were also recorded following the stimulation of a single AS with an increasing amplitude ramp. As can be seen in **Figure 5, B**, the amplitude of the positive peak was 1.1, 2.1, 3.9, 6.1, and 6.3 μV for 100, 200, 300, 400, and 500 μA respectively with nearly a 6-fold increase (**Figure 5, C**). Significant activation was visible with AS2, 4, and 5, while no activation was visible with AS 1 and 6. AS 2, which clearly showed the highest eCAP activity, was also the only AS which produced measurable physiological changes.

**Figure 5.**
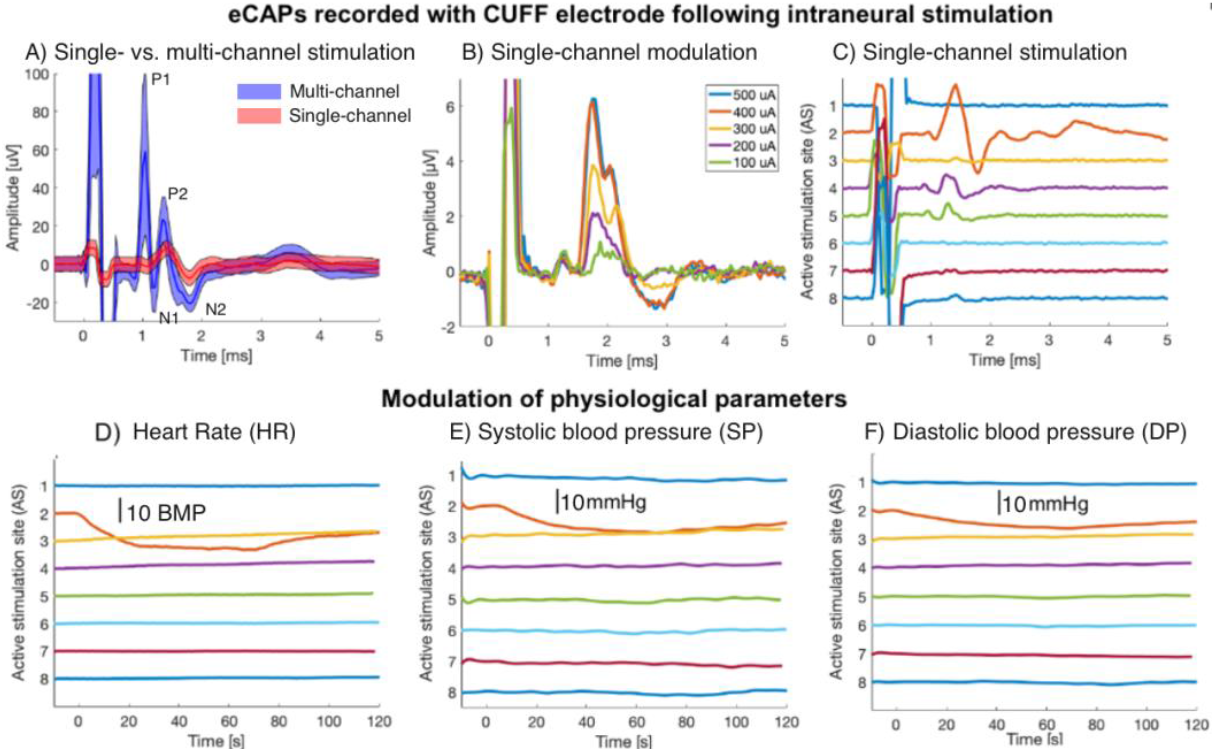
eCAP activity recorded CUFF electrodes and following neural stimulation through single-channel and multi-channel stimulation. A) Stimulation trigger is visible on the left, positive peaks (P1 and P2) and negative peaks (N1 and N2) are indicated. B) Activation of AB (P1), Ay (P2), and C (N1 and N2) fibers after the stimulation through intraneural stimulation with different amplitudes. C) eCAP activation following single-channel stimulation. D-F) Changes of physiological parameters (HR, DP, and SP) following single-channel AS stimulation.

### D. Physiological signal variation upon VNS

HR, DP, and SP were measured during single-channel stimulation. As shown in **Figure 5, D, E**, and **F**, a clear physiological response was observed only when stimulated using AS 2. The stimulation of AS2 reduced the HR by 13 beats per minute, while SP and DP decreased by 8.6 and 6.1 mmHg, respectively. No clear reduction was seen with any of the other AS. The results show that it was possible to elicit physiologically relevant responses using a single AS and only from that single AS. It is plausible that porcine VN structure can impact on the hemodynamic response to current injection.

## IV. DISCUSSION

### 1) Electrode design and development

Variations in the diameters of the nerve sections indicated that the minimum and maximum nerve diameter did not change significantly compared to previous anatomical vagus nerve studies where up to 31.7 % of variations occurred [16]. Stakenborg 2020 et al. reported an average nerve diameter of 2.9 mm where we found an average diameter of 2.64 mm. In the present study, only one animal was analyzed, making it difficult to draw further conclusions [17]. Stakenborg et al. reported 46 (std +/- 10) fascicles for the VN of animals of the same age and weight. With 37.2 (std +/-1.94), our measurements for the RVN are therefore close to the lower range of previously reported results. Since the nerve diameter we found for the RVN and LVN did not vary significantly, the number of fascicles was not related to the nerve dimension. In the present study two nerve samples were obtained. Inter-animal anatomical variation has therefore to be considered. Other studies performed a histological analysis corresponding to the mid-level of the cricoid reported by the present work. A std of 0.45 mm and 0.37 mm for the RVN max. and min. were reported respectively with a sample size of 6 nerves [18]. The importance of the here presented data is hence given, despite the small sample size. The nerve area of the LVN and RVN were similar with a difference of 0.3 *mm*^2^. Interestingly, the fascicle area was similar as well, indicating that the LVN with a similar diameter to the RVN contained more connective tissue (Table **2**). We have designed the electrode based on our histological data. AS distribution was based on the average nerve diameter. AS area (i.e., 0.005 mm2 corresponding to an AS ø of 80 μm) was smaller than the average LVN and RVN fascicle area of 1.2 and 1.6 mm2 respectively. The activation of single fascicle groups through intraneural stimulation was therefore given according to Oddo et al. 2016 [2].

### B. Electrode characterization

The measured AS diameter was increased compared to the designed AS. We believe that the precision of the AS is within an acceptable range. Improvements could be made by adapting the RIE process, and decreasing the etching time or adapting the electrode design. The ASH of the central and lateral devices varied within an acceptable range. TH and PH varied insignificantly. We could confirm that the photolithographic process of metallization is within range. The expected total height of the sputtered metal layers was Ti/Pt/IrOx 725 nm. The ASH was, on average, measured to be 35 nm higher than expected, whereas the TH was measured to be 0.7 nm higher than expected. Both values were in range according to the precision of the sputtering device (5 % according to the manufacturer). The ASH was 4.8 % greater than expected. However, a great variation between the ASH and the TH could be noticed. Since we measured central and lateral electrodes, we assume that the difference was caused due to the shape and dimension of the AS and track structure. We believe that the difference between the lateral and central devices could be neglected, which shows a uniform sputtering over the wafer. The difference in PH (measured and expected) was measured to be 2.5 % which was within range and was not expected to negatively influence conductive or mechanical properties. Higher precision may improve the development process, but not necessarily the quality of the intraneural stimulation. Hence, it is not necessary to focus on improving the development process. The here presented development method is valid and applicable for the design of a wide range of IE. The electrochemical analysis showed low impedance, optimal for the stimulation of the VN. Cogan et al. 2009 reported an impedance of 31.5 kΩ for 300 nm activated iridium oxide (AIROF) AS with an AS diameter of 50 μm on a 15 μm thick polyimide substrate [12]. Here we measured 12.4 kΩ for SIROF AS with 400 nm of IrOx on a IE made of similar material and substrate height with an AS diameter of 81.4 μm. The IE AS have a lower impedance due to a larger AS surface. Cogan et al. 2005 reported an average cCSC of 23 *mC*/*cm*^2^ for 50 μm diameter AS [8]. We found an average cCSC of 37.4 *mC*/*cm*^2^ for 80 μm diameter AS and were therefore within a reasonable range. All CV plots showed a counterclockwise inclination which can be seen in **Figure 4, B** and may indicate penetration of electrolyte (PBS) under the polyimide insulation [8]. In future development processes of IE adhesion promotion layers such as silicion carbide may be used to increase the adherence between polyimide and the metal structures [19]. The VTm data were averaged using a moving average filter, due to the low sampling rate of the oscilloscope. According to Cisnal et al. 2009, the maximum time window for V_a_ for the derivate of the VTm should be maximum of 10 μs [14]. The here reported time windows were 10 μs large, and therefore within the expected range. The MQ_inj_ of eight AS allowed to inject the maximum requested current amplitude of 500 μA at a pulse width of 200 ms. For the TIME, MQ_inj_ of 120 nC have been reported previously. Here we report a decrease of 21.4 nC (i.e. 17.8 %) in MQ_inj_ compared to the TIME. With a higher MQ_inj_ a wider range of stimulation could have been tested. However, 98.6 nC per AS was sufficient to intraneural activate VN fibers, as has been reported previously by Mc Callum 2017 et al. where charges between 4 and 25 nC have been applied using carbon nanofibers [20]. We assume that our IE had sufficient MQ_inj_, especially because single-channel stimulation with amplitudes between 400 and 500 μA did not show a further increase in eCAPs compared to stimulation with 300 and 400 μA. It is therefore suggested to develop neurostimulation systems such as presented here based on the stimulation patterns which should be applied.

### C. Neural selectivity

*In vivo* results confirmed previous studies showing that the selective modulation of the VN allows modulating the HR and BP [9] and showed that the here presented hardware could selectively modulate eCAP and physiological parameters. The delivered single-channel stimulation produced a significantly smaller evoked activity when compared to multi-channel stimulation. The presence of P1 and N1 only when all 8 AS are activated (**Figure 5, A**) suggests the recruitment of a much broader population of fibers in this configuration. Together, these suggest that single-channel stimulation is capable of activating a much smaller set of fibers, likely only those in proximity to the AS itself. Simulations using the present data would give a more detailed insight into this case. The development of a simulation model for the intraneural stimulation of the VN would be of great importance to the scientific community. The combination of eCAPs following neural stimulation and would allow validating such models, further improving the quality of neural stimulation in future experiments. Likewise, different levels of activation were observed when stimulated with different levels of intensity, with eCAPs increasing in size with progressively higher amplitudes up to 400 μA. This suggests that different levels of stimulation intensity can selectively activate sub portions of the fiber population that can be modulated by a single-channel stimulation. We thereby confirmed previously conducted studies showing this effect [7]. For future experiments, we suggest performing various combinations of AS stimulations to further increase neural selectivity. Furthermore, the difference between eCAPs recorded during single-channel stimulation, with a clear activation on AS 2, smaller activation on AS 4 and 5, and no clear activation on AS 1 and 6, confirms the ability of the electrode to activate different subsets of fibers. The P1 and N1 seen during multi-channel stimulation are not visible in any of the traces recorded, activating the AS one at the time. This suggests a non-linear summation of effects between the activation volumes of each individual AS where the total is greater than the sum of the parts. This suggests that activating multiple AS concurrently may allow for the modulation of fibers that cannot be activated by any single AS, increasing the flexibility of the stimulation. However, it still remains unclear what axon groups were activated due to eCAP measurement using cuff electrodes. To measure precise activation of axon groups more invasive electrodes would be necessary which may have a negative impact on IE signal propagation due to damage caused to the tissue. Lastly, it was possible to elicit physiologically relevant responses using a single AS. The fact that no other sites produced any visible response suggests that completely different population fibers can be modulated from two neighboring AS and confirms the high degree of selectivity of neuronal activation that can be obtained from the electrode.

## V. CONCLUSIONS

Here we developed neural stimulation hardware, including a novel IE and GUI and adapted an existing neurostimulator. We have shown that our approach to developing an IE based on a histological analysis allows for selective stimulation of the VN being a valid method. We showed that it was possible to selectively modulate the VN and resulting physiological functions. The present study represents an important base for the future development of neurostimulation hardware. The analysis of additional histological transections will allow us to specify the IE design. To increase selectivity further, the sitmulation between two AS could be performed. Experiments with an increased set of animals should be performed to show inter-animal reliability regarding neural selectivity. Multi-channel stimulation between AS and biomimetic stimulation may be applied to increase neural selectivity.

